# ASAP: A Machine-Learning Framework for Local Protein Properties

**DOI:** 10.1101/032532

**Authors:** Nadav Brandes, Dan Ofer, Michal Linial

**Affiliations:** Einstein Institute of Mathematics, The Hebrew University of Jerusalem, Jerusalem, Israel; Department of Biological Chemistry, The Hebrew University of Jerusalem, Jerusalem, Israel

**Keywords:** Feature extraction, cleavage sites, post translational modification, proteomics, machine learning

## Abstract

Determining residue level protein properties, such as the sites for post-translational modifications (PTMs) are vital to understanding proteins at all levels of function. Experimental methods are costly and time-consuming, thus high confidence predictions become essential for functional knowledge at a genomic scale. Traditional computational methods based on strict rules (e.g. regular expressions) fail to annotate sites that lack substantial similarity. Thus, Machine Learning (ML) methods become fundamental in annotating proteins with unknown function. We present ASAP (Amino-acid Sequence Annotation Prediction), a universal ML framework for residue-level predictions. ASAP extracts efficiently and fast large set of window-based features from raw sequences. The platform also supports easy integration of external features such as secondary structure or PSSM profiles. The features are then combined to train underlying ML classifiers. We present a detailed case study for ASAP that was used to train CleavePred, a state-of-the-art protein precursor cleavage sites predictor. Protein cleavage is a fundamental PTM shared by a wide variety of protein groups with minimal sequence similarity. Current computational methods have high false positive rates, making them suboptimal for this task. CleavePred has a simple Python API, and is freely accessible via a web-based application. The high performance of ASAP toward the task of precursor cleavage is suited for analyzing new proteomes at a genomic scale. The tool is attractive to protein design, mass spectrometry search engines and the discovery of new peptide hormones. In summary, we illustrate ASAP as an entry point for predicting PTMs. The approach and flexibility of the platform can easily be extended for additional residue specific tasks. ASAP and CleavePred source code available at https://github.com/ddofer/asap.

## 1. Background

Advances in protein functional prediction is an essential component for the interpretation of genomics data. Many protein properties that are of interest to the community can be localized to the level of individual amino acid (AA) residues in the primary sequence. These include post-translational modifications (PTM) sites such as proteolytic cleavage, as well as various local 1D structural states such as disorder and secondary structures.

The classic approach for functional annotation relies on sequence-similarity, augmented by multiple sequence alignments (e.g., HMM in Pfam (*1*)). Others resources such as PROSITE (*2*) and ELM (*3*) provide simple rules for proteins’ “signatures” (*4*). These methods suffer from high rates of false negatives and to a lesser extend false positives, making them suboptimal for genomic scale retrieval task. Proteins’ properties that cannot be represented in the form of a simple motif rely on sequence-based free models. Machine learning (ML) is a suitable approach for non-classical functional prediction challenges. Importantly, while ML methods benefit from the growth of sequences, the rule-based methods often fail to cope with sequence variability.

Several successful applications of residue level predictions using ML include secondary structure, disorder regions, functional families (*5*), PPI (*6*) and more. Despite the many ML classifiers, no generic framework (with easy to use API) is available for the task of predicting general residue-level properties. Initial effort in this direction includes ProFET project (*7*) that showed a success towards a broad range of biological classification challenges. In a similar line of thinking, we developed ASAP (Amino-acid Sequence Annotation Prediction) and demonstrate its usability toward predicting post-translationally cleavage sites from larger precursors. The ML classifier that is presented is CleavePred. It aims in correctly classify the cleavage sites for proproteins such as prohormones and neuropeptides. The processing proteases of proproteins in mammals belong to a diverse family of proteases called prohormone convertases (PCs). The unified rule for PCs is the presence of an arginine (R) or a lysine (K) at the first position N-terminal to the proteolytic sites. The predicting proteolytic cleavage sites occur throughout metazoan. The end products are active peptides and hormones that act in modulation in the endocrinic and neuronal systems.

In this short report we focus on the use of ASAP as a starting point for developing a high performing classifier that discriminate between any basic residue and the proteolytic sites. We discuss the results in view of the risk of model overfitting. The high precision obtained from ASAP and CleavPred suggest its usefulness in identifying likely candidates for experimental validation from unstudied sequenced genome.

## 2. Methods

### 2.1 ASAP pipeline

We are looking to solve the general problem of residue-level binary prediction (RLBP). Namely, predicting functional annotations for individual residues of a sequence. For example, we predict for each residue on a protein the score (0 or 1) for being a defined PTM sites (e.g., phosphorylation). To this end, we developed ASAP (Amino-acid Sequence Annotation Prediction), a universal Python framework for feature engineering and ML prediction. ASAP is completely generic, and can be easily applied to any task that involves binary predicting “local” properties of sequences, given a training dataset comprised of annotated sequences.

Applying ASAP to the case study of predicting cleavage sites in neuropeptide precursors, we created CleavePred, an ASAP-based model trained to solve the following RLBP task: for each residue along the precursor, predicting whether it is a cleavage site or not. ASAP providing the entire pipeline for feature extraction, transformation and model training. Both ASAP and CleavePred are free, open sourced (https://github.com/ddofer/asap), and come with a simple, flexible, and well-documented Python API. CleavePred is also accessible as a web-based application (http://protonet.cs.huji.ac.il/cleavepred). ASAP has a friendly tutorial (https://github.com/ddofer/asap/wiki/Getting-Started:-A-Basic-Tutorial).

The input of ASAP is a dataset comprised of annotated sequences in the “lf” (labeled file) format. Each annotation is a sequence of 0s and 1s for each position in the sequence. The input to ASAP is processed by the following step (Fig. 1): (i) Extraction of fixed-length windows. Each window is a sample in the training dataset. (ii) Windows may be filtered by various rules such as extraction only of sites centered on a K or R (in the case of CleavePred application). They may also be filtered by additional criteria, such as similarity to other windows. (iii) Sequence-based features are extracted for each window, creating feature vectors. (iv) The set of features are fed to a ML model for (v) training or prediction.

**Fig. 1.**
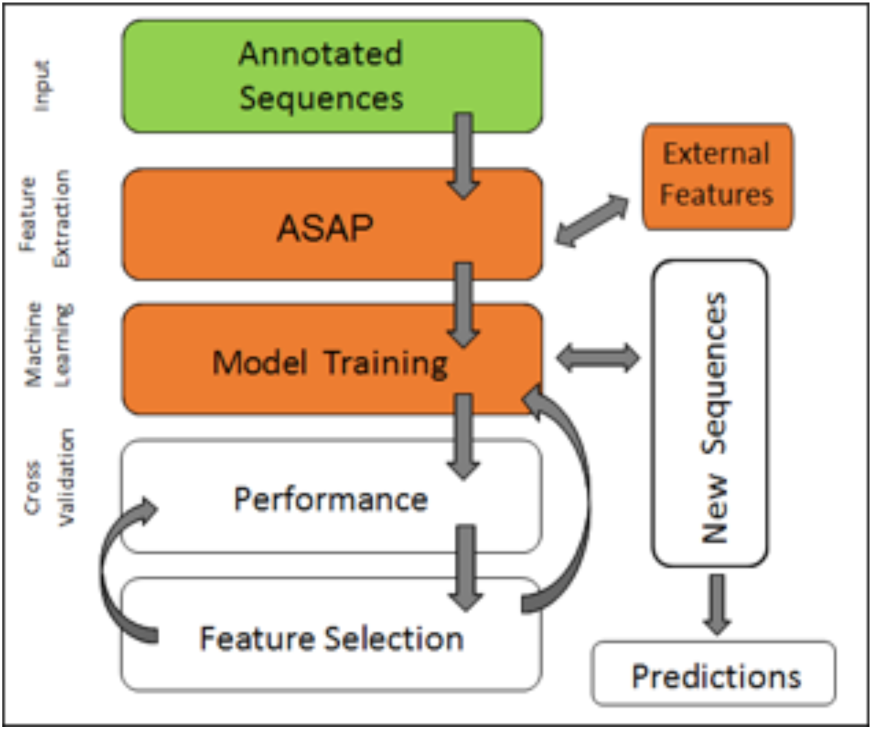
*ASAP, a feature extraction step in the prediction data workflow*

### 2.2 Feature extraction

Protein sequences are scanned, and fixed length windows are extracted, for each position or just for candidate sites (e.g. positions containing K or R in the case of CleavePred). Windows extending beyond the end of the sequence are padded with a dummy variable. A fixed length vector of features is extracted for each window. Additional features may be added from external predictors, notably 2D structure, solvent accessibility and disorder, via the SCRATCH (*8*) and DISOPRED3 toolkits (*9*), respectively.

### 2.3 ASAP supported features

ASAP supports multiple categories of features that are easily extendable. ASAP creates internally all features with the exception of secondary structure, solvent accessibility, disorder, PSSM, and PSSM entropy. We refer the readers to the API for in-depth details. ASAP includes freely available code to parse these features. Positions that exceeded the protein’s sequence were marked with a dummy variable, using one-hot-encoding (OHE). The features are groups according to their source and methodologies for feature engineering.

#### 2.3.1 Externally generated features

ASAP supports (optional) externally created features, including predictions made according to primary sequences. We currently support PSSM profiles, and predictions doe secondary structure, solvent accessibility and disorder. Our PSSM profiles were generated using SCRATCH’s ProfilPRO (V1.0) (*8*). Secondary structure (3 state resolution) and discretized solvent accessibility (buried or exposed) were predicted using SSpro and ACCpro. Discretized disorder predictions are obtained using DISOPRED3 (*9*)).

#### 2.3.2 Positional Features

These are properties relating to each individual position in a sequence. Discrete properties are encoded using OHE. These features are: AAs, Secondary structure, Disorder, AA Charge (±1 or 0), PSSM (frequency of the AA in the profile at a defined position) and Solvent accessibility. Additional engineered features include Reduced AA alphabets and PSSM entropy. The former is a lower dimensional representation of the AA alphabet, where “similar letters” are grouped together. We used a variant of 15 letters. The PSSM entropy can be seen as a measure of divergence of a position from a background or a uniform distribution. The more conserved a position is, the higher its entropy. The conservation score was calculated using the formula for relative entropy. The background for the CleavePred task is the AA frequencies in naturally occurring protein sequences from vertebrates.

#### 2.3.2 Contextual Aggregated Features

We applied the intuition accordingly local protein regions surrounding a site might have distinct aggregate properties. For the various positional features, we extracted aggregated averages over multiple positions. e.g. the “left” and “right” halves of the sequence.

For the PSSM entropy feature, we defined additional segments: (i) From the beginning of the window until position 4 residues (n-terminal-wise) prior to a potential cleavage site. (ii) The 4 residues prior to the putative cleavage site and the site itself. (iii) The remaining positions after the cleavage site (c-terminal-wise). We anticipated that a cleavage site would be more conserved as a whole than a random occurrence of flanking K|Rs’.

For feature defined as Disorder we used the naïve predictor FoldIndex that relies on hydrophobic potential and net charges (*10*) and TOP-IDP methods (*11*).

We will only mention briefly several groups of features that also are extracted by ASAP. Several of these features were extracted by ProFET (*7*).

#### 2.3.3 Motif Features

We combined the classic, “motif” based approach, as a regular expression when available. Specifically we have used a Cysteine spacer motifs (*12*) and dibasic sites (*13*).

#### 2.3.4 Quantitative Biophysical Features

Basic global features relating to biophysical properties of the window section were also gathered. The importance of these properties to validating proteins functions has been previously validated (*14*). These include feature such as Molecular weight, pH(I) but also the relative frequency of biophysically related group of AAs.

#### 2.3.6. AA scales based features

AA propensity scales can be used to represent the protein sequence as a time series, typically using sliding windows of different sizes and to extract additional features. We used maximally independent derived scales (*15*). It includes the averages for sequence segments and of a window as a positional feature. The default is a window size=4.

### 2.4. Datasets for CleavePred

Two Datasets of proteolytic cleavage were used: (i) NeuroPred dataset (*13*). (ii) The UniprotKB/Swiss-Prot, retried are all sequences that are manually annotated as having “cleavage on a pair of dibasic residues” and sequences carrying the feature “propeptide” / “peptide”. The datasets were filtered internally in two sequential steps (i) CD-HIT at a 65% similarity threshold; (ii) checking that no two proteins from different sets had >40% similarity. Signal peptides were removed.

### 2.5 CleavePred ML algorithms

We tested different models supported by scikit-learn (*16*). The final model used by CleavePred is an ensemble majority voting classifier (provided by the mlxtend package), using an SVM with a radial basis function kernel (C=3.798), a Random forest classifier and a logistic regression classifier. For each independent tested fold, features were filtered for zero variance, and ANOVA F-value (false discovery rate of q<0.1).

### 2.6 Models’ evaluation and testing

We trained “simple” and “advance” models for CleavePred. The simple model uses only sequence-based features, the advanced models also takes advantage of data obtained external tools (see 2.3). The two models were trained on NeuroPred’s dataset that contained, after reduced redundancy (see 2.4) 238 sequences. Of these ASAP extracted 6002 relevant windows (of basic AA residues) from which 4802 windows comprised the final training set. Evaluation was performed using a kfold procedure with 10 folds, were features were selected for each fold independently, based only on all training data. The models that were trained on NeuroPred’s dataset were then tested on the UniProt’s dataset. The latter contains 327 sequences after filtration with ASAP reports on 4325 windows and 3455 non-redundant ones.

The simple and advanced modes included 657 and 1352 features, respectively. After feature selection step, the number of features in the trained models was reduced to 482 and 960, respectively.

## 3. Results

We applied ASAP to the task of predicting cleavage sites (CleavePred). We trained and estimated our models on a specialized archive and database for neuropeptides, calls NeuroPred. We used a cross-validation (CV) procedure of 10 folds (Table 1). The results show the higher performance of the advanced model with respect to the simple mode. We then tested our trained models on UniProt’s dataset, and compared our performance to two competitor algorithms (Mammals and KM, Table 1). For our analysis, we insured that the training and test datasets are disjoint. Several conclusions can be drawn: (i) Our models (simple and advanced, test) are overall better and are superior in all measures. (ii) Maximal improvement is at the measure of precision (from 45–51% to 79%). (iii) The performance for the test set (UniProt) is lower in view of CV analysis (Table 1), as expected. However, Specificity remained extremely high (96%).

We further extracted the set of features that maximally contributed to high performance of our models (not shown). The top features allowed interesting biological interpretation that is beyond the scope of this abstract.

## 4. Discussion

In this study we present ASAP, a universal generic yet modular workflow for function prediction. The ASAP is useful as a platform (and an simple API) allowing an extensive analysis of new genomes and sequences that had not previously seen. This generic powerful framework can be applied to any (binary) residue-level problem (RLBP). In our tutorial, https://github.com/ddofer/asap/wiki/Getting-Started:-A-Basic-Tutorial, we demonstrate the usability of ASAP in approaching biological problems and obtain non-trivial results ASAP (i.e., in minutes). While there is always important fine-tuning phase and parameter optimization, we suggest using ASAP naively for a wide range of binary prediction tasks.

An important component is ASAP is the implementation of a combined approach. We combined naive features, basic feature engineering (e.g. aggregated features), and simple “rule based” approach (*i.e*., the canonical known motif), in a way that can easily be done for multiple tasks. This combined approach outperformed the state-of-the-art results substantially. Our approach also supports integration of external properties such as predicted secondary structure. This provides superior performance to either method alone.

Finally, we presented the power of ASAP towards a specific challenge of prohormones cleavage (CleavePred). For this task, we tested a more challenging training and validation set and reported the results on a novel, previously unseen test set (Table 2). We attribute the supreme performance and high confidence of our results to the engineered features that are the heart of ASAP, and the availability of high quality validated set of examples. CleavePred is extremely fast, making it suitable for whole-genome scanning. Due to the high cost of experimentally pursuing false-positives, the high precision of CleavePred allows focusing on only few high confidence candidates for further validation. CleavePred is accessible at http://protonet.cs.huji.ac.il/cleavepred.

**Table 2:** Performance of cross validation (CV) and Test sets

|  | **Simple**<br>**CV (%)** | **Advance**<br>**CV (%)** | **Simple**<br>**Test (%)** | **Advance**<br>**Test (%)** | **KM**<br>**Test (%)** | **Mammals**<br>**Test (%)** |
| --- | --- | --- | --- | --- | --- | --- |
| AUC | 88.17 | 89.08 | 80.42 | 76.75 | 75.94 | 81.94 |
| Accuracy | 93.48 | 94.40 | 89.87 | 88.68 | 72.45 | 77.81 |
| Sensitivity | 80.28 | 81.17 | 64.98 | 57.23 | 82.22 | 70.00 |
| Precision | 79.97 | 84.06 | 79.13 | 78.69 | 45.16 | 51.96 |
| Specificity | 96.07 | 96.99 | 95.87 | 96.26 | 69.46 | 80.20 |
| F1 Score | 80.13 | 82.59 | 71.36 | 66.26 | 58.30 | 59.64 |

